# Translesion DNA synthesis-driven mutagenesis in very early embryogenesis of fast cleaving embryos

**DOI:** 10.1101/2020.11.28.401471

**Authors:** Elena Lo Furno, Isabelle Busseau, Claudio Lorenzi, Cima Saghira, Matt C Danzi, Stephan Zuchner, Domenico Maiorano

## Abstract

In early embryogenesis of fast cleaving embryos DNA synthesis is short and surveillance mechanisms preserving genome integrity are inefficient implying the possible generation of mutations. We have analyzed mutagenesis in *Xenopus laevis* and *Drosophila melanogaster* early embryos. We report the occurrence of a high mutation rate in *Xenopus* and show that it is dependent upon the translesion DNA synthesis (TLS) master regulator Rad18. Unexpectedly, we observed a homology-directed repair contribution of Rad18 in reducing the mutation load. Genetic invalidation of TLS in the pre-blastoderm *Drosophila* embryo resulted in reduction of both the hatching rate and Single Nucleotide Variations on specific chromosome regions in adult flies. Altogether, these findings indicate that during very early *Xenopus* and *Drosophila* embryos TLS strongly contributes to the high mutation rate. This may constitute a previously unforeseen source of genetic diversity contributing to the polymorphisms of each individual with implications for genome evolution and species adaptation.

## Introduction

Very early embryogenesis of fast cleaving embryos is characterized by unusually contracted cell cycles, made of a periodic and synchronous succession of DNA synthesis (S-phase) and mitosis with virtually absent gap phases. S-phase length is dramatically short (15 minutes in *Xenopus* and only 4 minutes in *Drosophila*) and feedback mechanisms controlling genome integrity (checkpoints) are largely repressed, as there is no time to slow down the cell cycle (*1*) for review, and references therein). These include the ATR-dependent checkpoint that monitors replication fork progression (*2*)for review). This checkpoint is activated close to the midblastula transition (MBT) in concomitance with activation of zygotic transcription (*3*–*5*). Experiments in *Xenopus* have shown that checkpoint activation is sensitive to the DNA-to-cytoplasmic (N/C) ratio, since it can be triggered by artificially increasing the amount of DNA in the embryo over a threshold level, a situation that mimics the increase in DNA content reached close to the MBT (*6*). Previous observations in *Caenorhabditis elegans* (*7*–*9*) and more recently in *Xenopus laevis* (*10*) have implicated the translesion DNA synthesis (TLS) branch of DNA damage tolerance in silencing the DNA damage checkpoint. In *Xenopus* cleavage-stage embryos, constitutive recruitment of at least one Y-family TLS polymerase (Pol η) onto replication forks, driven by the TLS master regulator Rad18 (E3) ubiquitin ligase, minimizes replication fork stalling in front of UV lesions thereby limiting ssDNA production which is essential for replication checkpoint activation (*10*–*13*). This configuration is lost prior to MBT following a developmentally-regulated decline of Rad18 abundance (*10*).

TLS Pols have the unique capacity to replicate damaged DNA thanks to a catalytic site more open than that of replicative polymerases which can accommodate damaged bases. Because TLS Pols cannot discriminate the insertion of the correct nucleotide and lack proofreading activity, they can be highly mutagenic especially on undamaged templates (*14*)for review). Recruitment of Y-family TLS pols (ι, η, κ and Rev1) requires monoubiquitination of the replication fork-associated protein PCNA (PCNA^mUb^) by Rad18 (E3) and Rad6 (E2) ubiquitin ligases complex (*15*, *16*). Aside from its TLS function, Rad18 is also implicated in error-free homology-directed DNA repair (HDR) in response to both double strand breaks (DSBs) and interstrand cross-links (*17*–*21*). These functions are separable and lie in distinct domains of the Rad18 protein. The Rad18 TLS activity is confined to its ring finger domain (*22*), while the HDR activity mainly depends upon its zinc finger and ubiquitin binding domain (*17*, *18*). We have previously shown that in early *Xenopus* embryos PCNA is constitutively monoubiquitinated, irrespective of the presence of DNA damage (*10*). Whether TLS is active during the early embryonic cleavage stages is currently unclear. Previous work in *C. elegans* has shown that mutations in some TLS Pols do not influence global mutagenesis although a *polη* and *polκ* double mutant accumulate DNA deletions (*9*). In this work, we provide evidence for TLS-dependent mutagenesis in early *Xenopus* and *Drosophila* embryos and show that in *Xenopus*, both Rad18 HDR activity and the mismatch repair system (MMR) alleviate mutagenesis, thus reducing the mutation load.

## Materials and Methods

Experiments with *Xenopus* were performed in accordance with current institutional and national regulations approved by the Minister of Research under supervision of the Departmental Direction of Population Protection (DDPP). *Xenopus* embryos were prepared by *in vitro* fertilization as previously described (*10*). Two-cell stage embryos were microinjected in the animal pole using a Nanoject auto oocyte injector under a stereomicroscope (2 injections of 9 nL^−1^ in one blastomer). Each embryo was injected with 12 ng (pre-MBT) or 72 ng (post-MBT) of supercoiled plasmid, undamaged or irradiated with 200 J/m^2^ of UV-C with a UV-Stratalinker, and/or 5 ng of RAD18 mRNAs. Embryos were collected at 16- or 32-cell stage, according to Nieuwkoop and Faber normal tables, snap frozen in liquid nitrogen and stored at −80 °C.

*Drosophila melanogaster* stocks were maintained and experiments were performed following standard procedures on standard cornmeal-yeast medium, inside a thermostatic room at 25 °C with alternating light and dark for an equal amount of hours per day. The following stocks were from Bloomington *Drosophila* Stock Center : *dPolηExc2.15* (#57341), OreRmE (#25211). The latter was homogeneized through serial individual backcrosses for nine generations. *dPolη^12^* was a kind gift of Benjamin Loppin (*23*). Balancer stocks were from our laboratory. Quantifications of hatching rates (eggs to larvae) were determined as previously described (*24*). Hatch rate is the ratio of hatched eggs to total eggs laid expressed as a percentage. Two hundred or more embryos were scored twice per genotype. In addition, hatching of adult flies was estimated by calculating the percentage of larvae (counted 2-2,5 days after fertilization) developed to mature flies (counted 10 days after fertilization).

### Plasmid DNAs

*lacZ*-containing plasmid (pEL1) was obtained by subcloning the *lac* operon from pBluescript into the *Spe*I-*Kpn*I restriction sites of pRU1103 vector, which contains full-length *lacZ*. pEL1 was transformed and amplified in *E. coli* and purified using a standard protocol (QIAGEN) at a temperature lower or equal to 12 °C to obtain a near 100 % supercoiled DNA, as previously described (*25*). This procedure greatly minimizes DNA damage and background mutations. pCS2-MLH1 plasmid was obtained by subcloning human MLH1 cDNA from pCEP9MLH1 (*72*)), into the *BamH*I-*Xho*I restriction sites of pCS2 vector. Rad18 wild-type, C28F and C207F mutant plasmids were previously described (*10*). The Rad18^C28FC207^ double mutant was generated by standard site-directed mutagenesis from the Rad18^C28F^ mutant plasmid.

### In vitro transcription

mRNA synthesis was performed with mMESSAGE mMACHINE kit SP6® (AM1340, Thermofisher). mRNAs were recovered by phenol-chloroform extraction and isopropanol precipitation. Following centrifugation and ethanol wash, mRNAs were dissolved in 20 μL^−1^ of RNase-free water. mRNAs quality was checked by formaldehyde gel electrophoresis.

### Xenopus embryos and eggs protein extracts

An average of 20 embryos were lysed in Xb buffer (5 μL^−1^ of buffer per embryo; 100 mM KCl, 0.1 mM CaCl_2_, 1 mM MgCl_2_, 50 mM sucrose, 10 mM HEPES pH 7.7) supplemented with cytochalasin (10 μg mL^−1^), phosphatases (PhosSTOP 1×,) and proteases inhibitors (5 μg mL^−1^ Leupeptin, Pepstatin A and Aprotinin). After 10 min centrifugation at maximum speed in a benchtop centrifuge at 4 °C, cytoplasmic fraction was recovered, neutralized in an equal volume of Laemmli buffer 2× and boiled at 95 °C for 5 min. Embryos lysates were loaded on precast gradient gels (4-12 %, NuPAGE, Invitrogen). Gels were transferred to a nitrocellulose membrane for western blotting and incubated with the indicated antibodies. Interphasic *Xenopus* egg extracts were prepared and used as previously described (*10*).

### Ribonucleotide incorporation assay

Upon thawing, *Xenopus* eggs extracts were supplemented with cycloheximide (250 μg mL^−1^) and an energy regeneration system (1 mM ATP, 2 mM MgCl_2_, 10 mM creatine kinase, 10 mM creatine phosphate). M13mp18 ssDNA was added as a template for DNA replication at the indicated concentrations in presence of α-;^32^PdCTP (3000Ci mmol^−1^, Perkin Elmer). At the indicated time points half of the samples were neutralized in 10 mM EDTA, 0,5 % SDS, 200 μg mL^−1^ Proteinase K and incubated at 52 ° C for 1 hour. Samples were treated with 0.3 M NaOH at 55°C for 2 hours to digest incorporated ribonucleotides in the plasmid and loaded on 5M urea 8% acrylamide gel TBE 0,5 X urea after formamide denaturation at 55 °C for 3 minutes. After migration, the gel was exposed to autoradiography.

### Plasmid DNA isolation from embryos

Frozen embryos were crushed in STOP MIX supplemented with fresh Proteinase K (600 μg μL^− 1^). Embryos were homogenized with a tip in this solution while thawing. Immediately after, proteins digestion at 37 °C for 1 hour, total DNA was extracted as described above by phenol-chlorophorm extraction and ethanol precipitation. Recovered DNA was digested with *Dpn*I to destroy unreplicated plasmids and subsequently purified with QIAGEN gel extraction kit.

### Somatic A6 cell culture

A6 epithelial cells were grown in modified Leibowitz L-15 medium containing 20 % sterile distilled water, 10 % foetal bovine serum and 100 U/ml penicillin/streptomycin at 25°C. A subcultivation ratio of 1:3 was employed. Cells were detached after a single wash with PBS by incubation with 0.25 % trypsin 0,03 % EDTA for 4 minutes at 37 °C. The day prior to transfection, 3 × 10 ^−6^ cells were seeded in 10 cm^2^ dishes. One day later, cells were transfected with 60 μL Lipofectamine 2000 (Thermo Fisher Scientific) and 24 μg of plasmid following the manufacturer’s recommendations.

### Plasmid recovery from A6 cells

Cells were harvested by trypsinization and centrifugation 48 hours post transfection. After washing cell pellet with PBS, cells were crushed in lysis buffer (10 mM Tris pH 8,0; 100 mM NaCl; 10 mM EDTA pH 8,0; 0,5 % SDS) supplemented with fresh Proteinase K (600 μg μL^−1^) by means of a tip (500 μL^−1^ per sample). Immediately after protein digestion at 37 °C, SDS was precipitated by adding half volume of saturated (6M) NaCl to each tube and centrifugation at 4°C for 10 minutes at 5000 rpm after 10 minutes incubation on ice. Total DNA was extracted from the supernatant as described above by phenol-chlorophorm extraction and ethanol precipitation. Recovered DNA was digested with *Dpn*I restriction enzyme (NEB) to destroy unreplicated plasmids and subsequently purified with QIAGEN gel extraction kit.

### White/blue colonies selection and mutation frequency

DNA extracted from embryos was transformed in electrocompetent indicator bacteria (MBM7070 strain bearing an amber mutation in the *lacZ* gene) for white/blue screening and plated on selective petri dishes (40 μg mL^−1^ Xgal,; 200 μM IPTG). Over one thousand colonies were scored at least for each condition in each replicate. Plasmid DNA was isolated from mutant clones using a standard protocol (QIAGEN). After paired-end Sanger sequencing, polymorphisms were filtered for sequencing quality > 30 and analyzed on both strands using Geneious or Snapgene softwares. Mutation rates were estimated from the proportion of blue colonies observed (P_0_). Before calculating the proportion of blue colonies observed (P_0_), the basal percentage of white colonies prior to microinjection was subtracted from the percentage of white colonies in each experimental condition. The observed P_0_ was substituted for P_0_ to obtain the mutation rate (μ) using the following formula: μ=-ln(P_0_) and normalized to the number of cell cycles before embryo collection.

### Antibodies

The following antibodies were used: Gapdh (ab9484, Abcam); Pcna^mUb^ Lys 164 (13439, Cell Signaling Technology); PCNA (PC10, Sigma); XlRad18 (*10*); SMAUG (*27*); Tubulin (DM1A, Sigma), Mlh1 (ab14206 Abcam). The PC10 antibody cross-reacts with *Drosophila melanogaster* PCNA (*28*).

### DAPI staining of Drosophila embryos

Embryos collection (0-2 hours unless otherwise indicated) was carried out using standard techniques (*29*). Embryos were dechorionated in 50% bleach and fixed by shaking in a mixture of PFA 4% in PBS and heptane (1:1) for 30 minutes and the aqueous layer containing formaldehyde was removed. Embryos were devitellinised upon washing in methanol-heptane mixture (1:1) and conserved in methanol at −20 °C overnight and for up to a week. Embryos were rehydrated by sequential incubations of 10 minutes in Ethanol/PBS-T 7:3 (1X PBS + 0.1% Triton X-100), Ethanol/PBS-T 3:7 and PBS-T. Embryos were incubated for 30 minutes at room temperature in DAPI-PBS-T (1μg mL^−1^) in the dark and rinsed three times in PBS-T. The third wash was performed overnight with mild shaking on a wheel in the dark at 4 °C. Samples were mounted in coverslips using Vectashield. Images were acquired with a Zeiss Axiovert Apotome microscope at 5× using Coolsnap HQ CDD camera (Photometrics) and processed using Omero 5.2.0 software. *P*-values were obtained using a two-tailed, unpaired Student’s *t*-test.

### Drosophila embryo protein lysates preparation

60 females and 10 males were incubated together inside embryo’s collectors with embryo dishes for a certain number of hours according to the desired stage of embryos to be harvested. Collected embryos were gently rinsed off the medium with embryo collection buffer (Triton X-100 0,03 %; NaCl 68 mM). Embryos were removed from the medium using a brush and poured into a sieve (Falcon Cell Strainer 40 μM Nylon 352340). Harvested embryos were washed again and collected in a fresh tube (up to 50 μL^−1^ of embryos corresponding to 100 embryos). Laemmli 2X was added in ratio 1:1 in comparison to harvested volume and embryos were lysed by means of a pestle. After boiling embryo’s mush at 95 °C for 5 min, chorion residues were removed by centrifugation with a benchtop centrifuge (max speed) at room temperature. Protein concentration was estimated by Amido Black staining using BSA of known concentration as a reference.

### *Genomic DNA extraction from single flies* for Next Generation Illumina Sequencing

Each fly was crushed with a pestle in a 1,5 mL^−1^ tube containing 170 μL of extraction buffer (Tris-HCl pH 8.0; 50mM; EDTA 50mM; SDS 1 %). Proteinase K (555 μg μL^−1^) was added once the tissues had been completely grinded. After incubating 15 min at room temperature, cellular debris was removed twice by potassium acetate addition (0,83 M) and centrifugation with a bench-top centrifuge (max speed) at 4 °C for 10 min. DNA was isolated by double phenol/chloroform/isoamyl alcohol (25:24:1) extraction and ethanol precipitation overnight at −20 °C with glycogen (20 μg) followed by centrifugation at 4 °C. Precipitated DNA was washed with cold ethanol 70 %, dried at room temperature for 30 min and dissolved in water. The quality of extracted DNA (about 150 ng) was verified by agarose gel electrophoresis.

### Illumina Next Generation Sequencing

Two genomic DNA samples per condition (extracted from single heterozygous *dpolη^EXC215/+^* males either dPolη maternally-depleted or maternally-provided, see text) were sequenced by Illumina NGS. After library construction and shotgun, whole *Drosophila* genomes were paired-end sequenced and assembled as previously described (*30*). Data were filtered for Genotype Quality > 35 and Depth > 10 before sequences alignment against the *Drosophila* reference genome (Flybase release 6). Variant calling was done using Freebayes software version 0.9.20. Genotype ratio was not changed from recommended settings. Alignment was performed with BWA version 0.7.12-r1039. Variant calling was performed using Freebayes version 0.9.20 and then annotated with Ensembl VEP version 82. SNVs and Indels were then separated for downstream analyses. The threshold generally is above 33% to call an allele variant from the reference.

### Statistics

Statistical analysis was performed using the Prism software (version 8). Means were compared using analysis of one-way ANOVA. Post-hoc tests were performed with a two-tailed unpaired Student’s t test unless otherwise specified. Stars indicate significant differences * P< 0.05, ** P< 0.01, *** P <0.001, **** P <0.0001, “ns” denotes non-significant statistical test.

## Results

### High mutagenesis rate in the pre-MBT *Xenopus* embryo

We employed a classical *lacZ*-based reporter assay to measure mutagenesis in pre-MBT *Xenopus laevis* embryos. In this experimental procedure, a plasmid containing the whole 3 kb *lacZ* gene is microinjected in *in vitro* fertilized *Xenopus* embryos at the 2-cell stage (Figure 1A) and development is allowed to continue until before MBT (16-cell stage). Upon injection, plasmid DNAs form minichromosomes and replicate as episomes once per cell cycle with no sequence specificity (*31*, *32*). Total DNA is then extracted, purified and plasmid DNA is recovered in *E. coli* by transformation, since only plasmid DNA can transform bacteria (see Materials and Methods). Bacteria are plated on a chromogenic substrate (X-gal) to screen colonies for white or blue color. Wild-type *lacZ* produces active β-galactosidase which stains colonies in blue in the presence of X-gal and IPTG, while mutations generated in the *lacZ* gene that affect β-galactosidase activity will leave colonies colorless (white) or pale blue. A pre-MBT dose of supercoiled plasmid DNA (12 ng/embryo, Supplementary Figure S1A) was used in most of the experiments as previously described (*6*).

**Figure 1.**
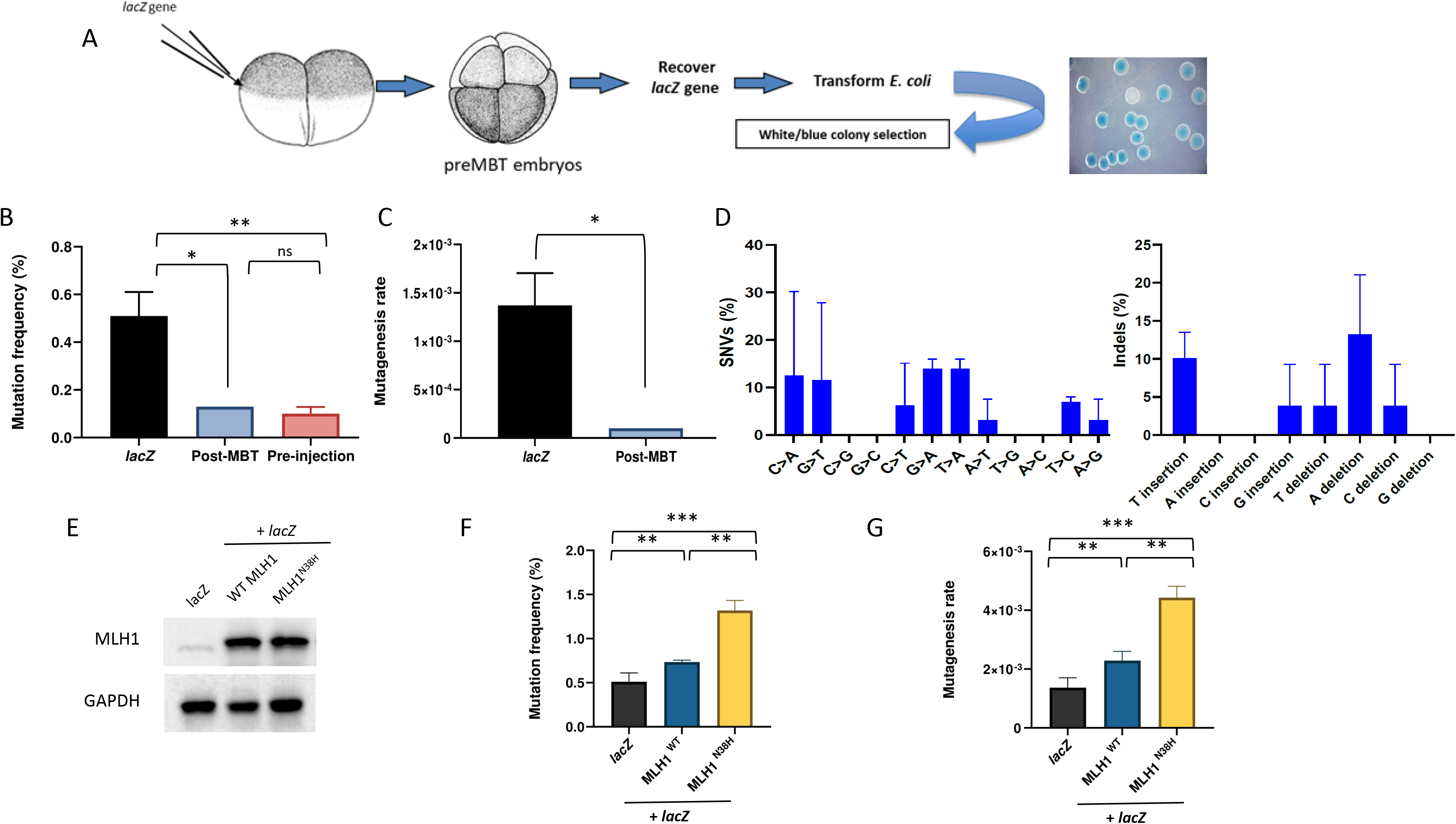
Pre-MBT *Xenopus* embryos accumulate polymorphisms and deletions. (A) Drawing of the experimental strategy adopted to analyze mutagenesis in *Xenopus laevis* embryos. 2-cell stage embryos are injected with a supercoiled plasmid containing *lacZ*-reporter gene (pEL1) and allowed to replicate for further 3 divisions. After embryos collection, plasmid DNA is extracted and transformed in *lacZ*-deficient bacteria for white/blue screening. (B) Mutation frequency expressed as percentage of white colonies in each condition. The mutation frequency of *lacZ* recovered from embryos injected with a post-MBT amount of plasmid DNA is also included as comparison. pre-MBT and post-MBT n=3, pre-injection: n=2. (C) Mutagenesis rate in the indicated different experimental conditions expressed as mutations per base pair/locus per generation (see Materials and Methods), normalized to the pre-injection background values, n=3. (D) Mutation spectra of the *lacZ* gene recovered from *Xenopus* pre-MBT embryos after Sanger sequencing, n=3. (E) Western blot of total protein extracts obtained from *Xenopus* embryos subjected to the indicated experimental conditions, n=2. (F) Mutation frequency and (G) mutagenesis rate of *lacZ* isolated from *Xenopus* embryos injected as indicated. *lacZ* n=3; Mlh1: n=2. Data are presented as means ± SD. n=2. Means were compared using unpaired Student’s t test.

Recovery of the *lacZ*-containing plasmid DNA isolated from pre-MBT embryos into *E. coli,* generated white colonies with a frequency of 0,5 %, compared to the non-injected plasmid (pre-injection, Figure 1B) or to the same plasmid transfected into *Xenopus* somatic cells (Supplementary Figure S1E). Accordingly, mutation rate was calculated by normalization to the number of cell cycles (see Materials and Methods) and estimated to be in the order of 10^−3^ (Figure 1C, *lacZ*). Importantly, the mutation rate dropped to a background level when embryos were injected with a post-MBT amount of plasmid DNA, a situation that increases the N/C ratio and induces a cell cycle delay (*6*). Analysis of mutations by DNA sequencing revealed the presence of both single nucleotides variations (SNVs) and unexpectedly large deletions ranging from 100 bp to 1,5 kb (Figure 1D and Supplementary Figure S1B). Mutations inspection on the *lacZ* gene showed that they are generally widespread over the entire sequence with no hotspots (Supplementary Figure S1B). Analysis of the mutation spectrum shows that most SNVs detected were C>A and C>T changes (Figure 1D). Another frequent signature was G>A transitions and T>A transversions, as well as nucleotides insertions and deletions. This mutation spectrum is close to that reported for TLS Pols on undamaged templates (*14*), in particular Polη and Polκ (*33*, *34*), although C>T transitions are also thought to be due to spontaneous deamination of 5-methyl cytosine to thymine.

The high frequency of base substitution and deletion prompted us to test the contribution of the mismatch repair system in the mutagenesis rate. For this, we overexpressed either wild-type or a catalytically inactive mutant (N38H) of Mlh1, a critical MMR component (*10*). We co-injected the *lacZ*-containing plasmid together with *in vitro-* transcribed MLH1 mRNAs to act as dominant negative by antagonizing the function of the endogenous protein (Figure 1E). While expression of Mlh1^WT^ only slightly increased the mutation rate, this latter was increased 2-fold upon expression of the Mlh1^N38H^ catalytically-inactive mutant (Figure 1E-G) suggesting that the MMR is functional and contributes to restrain mutagenesis. Altogether, these results show that the mutation spectrum observed in pre-MBT *Xenopus* embryos is similar to that expected for TLS Pols and that mutagenesis is restrained by the MMR system, suggesting that TLS Pols may actively contribute to mutagenesis in very early embryogenesis.

### Rad18-dependent mutagenesis in the early *Xenopus* embryos

We have previously shown that in the pre-MBT *Xenopus* embryo TLS may be constitutively primed at replication forks in absence of external DNA damage (*10*). To determine the possible contribution of TLS to the mutagenesis in *Xenopus* embryos, we made use of a Rad18 TLS-deficient mutant in a dominant negative assay as done for Mlh1 (Figure 2A). Mutagenesis was analyzed as described in the previous paragraph. Expression of the TLS-deficient Rad18^C28F^ mutant strongly reduced both the frequency of white colonies and the mutagenesis rate of about 100-fold compared to injection of either Rad18^WT^ or *lacZ* alone (Figure 2B-C). In contrast, Rad18^WT^ overexpression did not alter the mutagenesis rate compared to embryos injected with *lacZ* plasmid only, although it generated a different mutational spectrum, consisting of T>A transversions, C and G insertions, and remarkably no large deletions (Figure 2E and Supplementary Figure S2A). T>A transversions were reported to be significantly decreased in mice bearing the *pcna^K164R^* mutation that cannot support PCNA^mUb^ (*35*), suggesting that this signature is Rad18 TLS activity-dependent. Compared to Rad18^WT^, the residual mutagenesis observed in embryos injected with the Rad18^C28F^ mutant showed a drastically reduced frequency of T>A transversion as well as C and T insertions, a TLS Polη and Polκ signature, suggesting that these mutations are PCNA^mUb^-dependent, while the frequency of T>C transitions increased. These latter mutations are consistent with a Rev1 signature, a TLS Pol that can also be recruited independently of PCNA^mUb^ (*36*, *37*).

**Figure 2.**
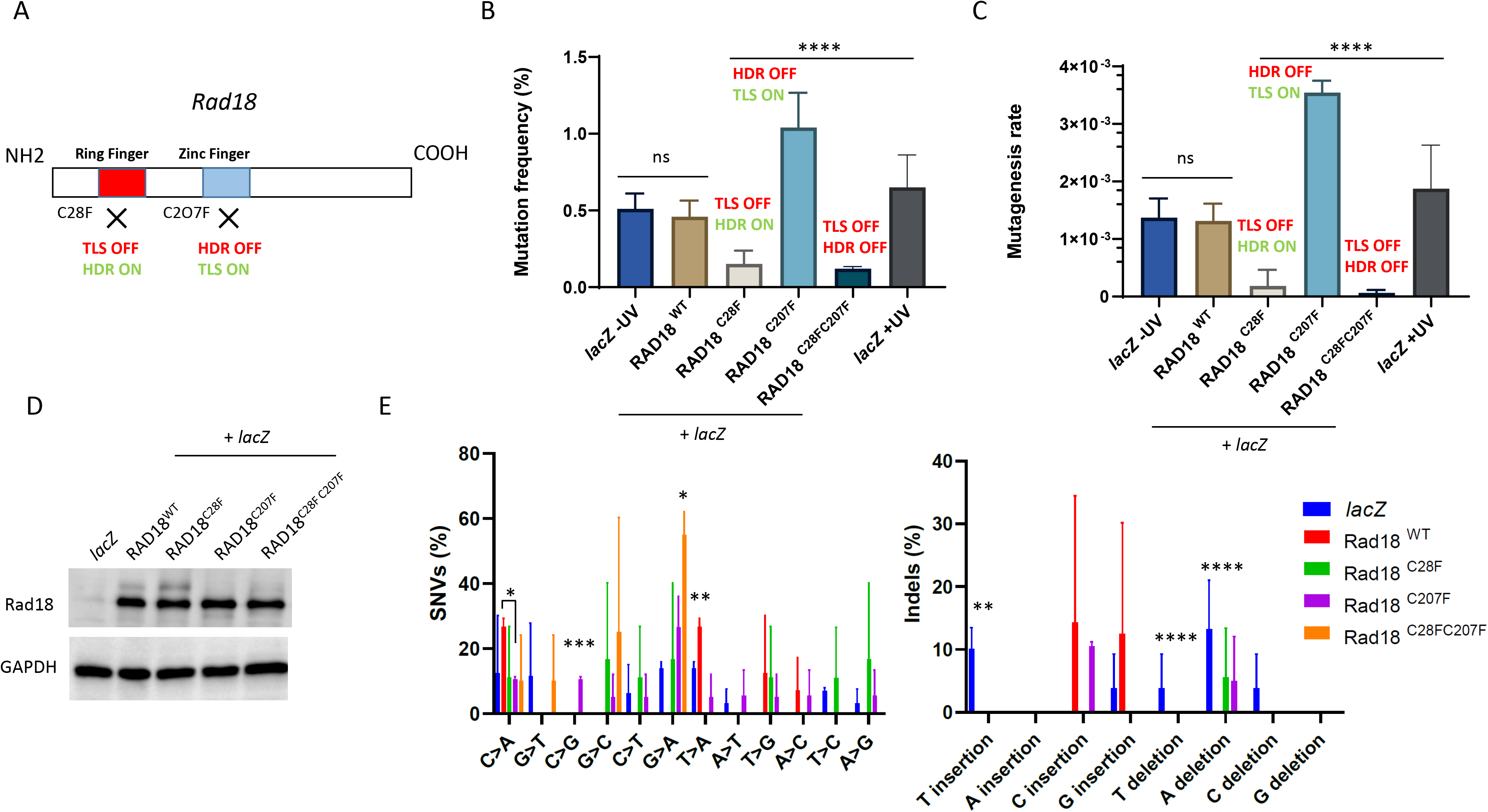
Differential contribution of Rad18 to mutagenesis in pre-MBT *Xenopus* embryos. (A) Schematic illustration of Rad18 domains in DNA damage tolerance and repair. TLS depends on the ring finger domain, while HDR is dependent on the zinc finger domain. The C28F mutation knocks out TLS activity (TLS OFF) while the C207F mutation knocks out HDR activity (HDR OFF). (B) Mutation frequency and (C) mutagenesis rate (measured as described in Figure 1) of *lacZ* recovered from embryos co-injected with the indicated RAD18 mRNAs, or *lacZ* injected alone. The mutation frequency of *lacZ* recovered from embryos injected with a post-MBT amount of plasmid DNA (post-MBT) is also included as comparison. RAD18^WT^, RAD18^C28F^ n=3; RAD18^C207F^, RAD18^C28FC207F^ n=2. (D) Western blot of total protein extracts obtained from *Xenopus* embryos subjected to the indicated experimental conditions, n=3. (E) Mutation spectrum of *lacZ* recovered from embryos injected with the indicated RAD18 variants, or *lacZ* injected alone. Data are presented as means ± SD. Means were compared using analysis of one-way ANOVA, followed by unpaired Student’s t test.

We also tested the effect of expressing the homology-directed repair (HDR)-deficient, TLS-proficient, Rad18^C207F^mutant, which is predicted to behave as Rad18^WT^ (Figure 2A). Unexpectedly, however, expression of this mutant increased the number of white colonies of 2-fold compared to Rad18^WT^ or *lacZ* alone, and the mutagenesis rate increased accordingly (Figure 2B-C) notwithstanding a similar expression level (Figure 2D). Compared to Rad18^WT^, expression of the Rad18^C207F^ mutant produced a reduction in both T>A transversions (Figure 2E) and generated large deletions (Supplementary Figure S2C and see below). The significant increase in single nucleotide substitutions generated by this mutant is consistent with the occurrence of unproofed mutations generated by its TLS activity. As expected, expression of the Rad18^C28FC207F^ double mutant produced a mutation burden similar to that of the TLS-deficient Rad18^C28F^ mutant, strongly suggesting that the mutagenesis restricted by the Rad18 HDR activity is TLS-dependent. Expression of this mutant increased G>A transitions and also produced large deletions (Figure 2E and Supplementary Figure S2D). In parallel, we analyzed mutagenesis when TLS is normally activated by UV irradiation and observed a very modest increase. The mutation spectrum was similar to that of the –UV condition and showed an increase in T insertions, as expected, which corresponds to TLS Polη and Polκ mutational signature (*33*)(Figure 2E and Supplementary Figure S1C-D), as well as disappearance of C>G transversions and reduction of C>T transitions. The modest increase in UV-induced mutagenesis is expected if TLS is constitutively activated, and is consistent with an error-free bypass of UV lesions by TLS Polη. Interestingly, no large deletions were detected (Supplementary Figure S1C and see discussion). Collectively, these results show that mutagenesis in the pre-MBT *Xenopus* embryo is Rad18-dependent and that, unexpectedly, the extent of TLS-dependent mutagenesis is alleviated by the error-free Rad18-dependent HDR activity.

### Reduced hatching rate in *dpolη* maternally-deprived flies

In the aim to assess whether TLS-dependent mutagenesis is a general feature of fast cleaving embryos, and to obtain genetic evidence for this process, we turned to *Drosophila melanogaster*, a more genetically amenable system compared to allotetraploid *Xenopus*. First, we wished to establish whether developmental regulation of PCNA^mUb^ also occurs during *Drosophila* embryogenesis. Similar to *Xenopus*, *Drosophila* early development occurs through a rapid and synchronous series of embryonic cleavages before activation of zygotic transcription (MBT, Figure 3A)(*38*). Total protein extracts were prepared from *Drosophila* embryos before and after MBT and both total PCNA and PCNA^mUb^ levels were analyzed by western blot with specific antibodies (see Materials and Methods and Supplementary Figure S3A). Figure 3B-C shows that similar to what previously observed in *Xenopus* (*10*), PCNA^mUb^ is detectable in pre-MBT *Drosophila* embryos (0-2 hours) and declines at later stages (3-5 hours, post-MBT). The developmental stage where a decline in PCNA^mUb^ is observed coincided with that of Smaug, a mRNA polyadenylation factor destabilized just after MBT (*39*). We could not probe Rad18 expression since a *Drosophila* ortholog could not be found, neither by sequence homology, nor by structure-specific alignments (Busseau, Lo Furno, Bourbon, and Maiorano, unpublished). Altogether, these observations suggest that in pre-MBT *Drosophila* embryos TLS may be constitutively primed. In line with this conclusion, previous observations have shown that *Drosophila* Polη (dPolη) is highly expressed in pre-MBT embryos, localizes into interphase nuclei, similar to what previously observed in *Xenopus* (*10*), while *dpol*η mutant embryos are sensitive to UV-irradiation (*40*).

**Figure 3.**
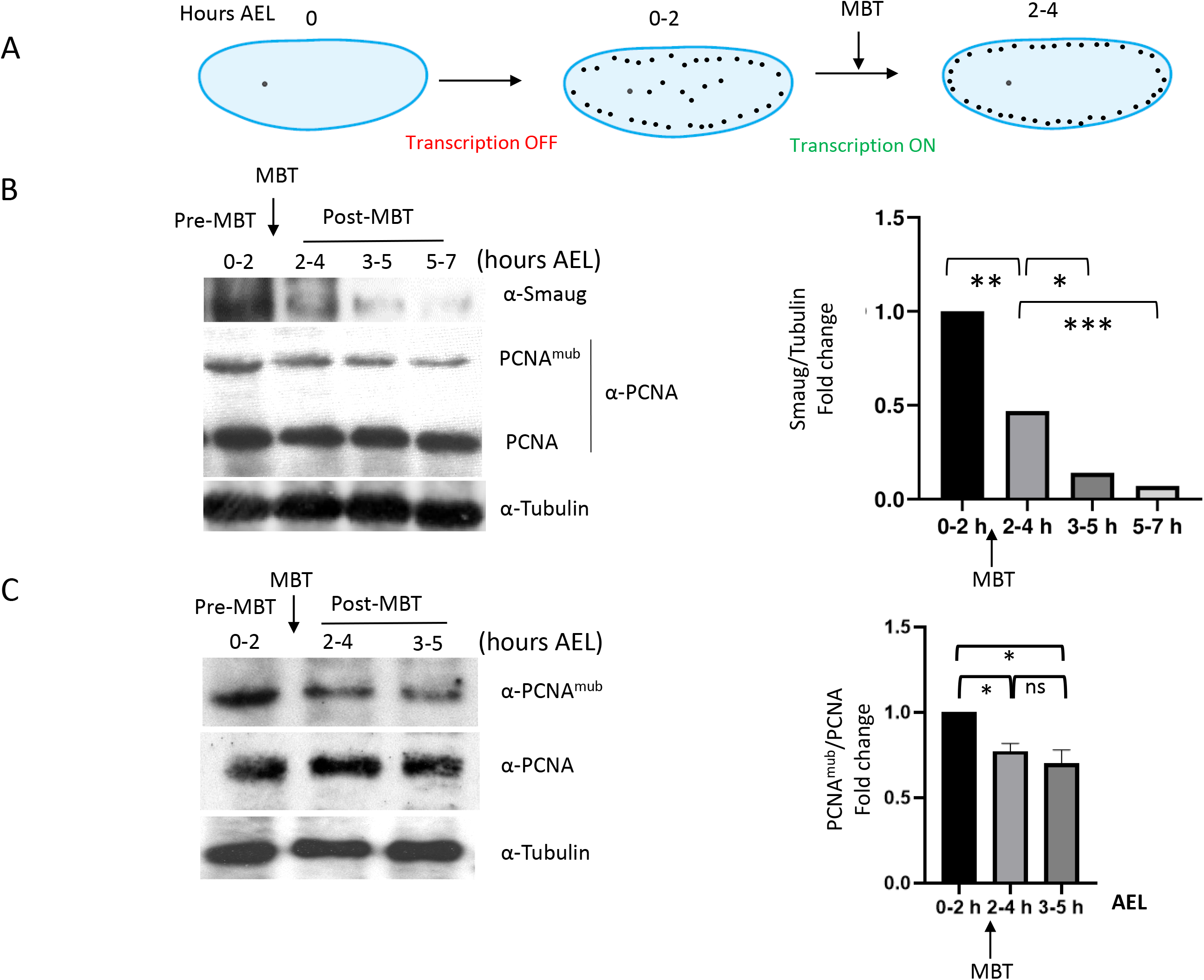
Developmental regulation of PCNA^mUb^ in *Drosophila*. (A) Simplified scheme of early *Drosophila melanogaster* embryogenesis. Embryos were collected at different hours after eggs laying (AEL), pre-MBT (0-2h), MBT (2-4h) and post-MBT (3-5 h and 5-7h). (B-C, left panels) Western blot of total *Drosophila* embryos extracts analyzed at the indicated times with the indicated antibodies. Right panels, quantification of western blot panels. Data are presented as means ± SD. Means were compared using unpaired Student’s t test.

In *Drosophila*, the presence of only three Y-family TLS Pols has been reported, since a Polκ ortholog could not be identified (*41*). In order to gain insight into whether TLS is also active in pre-MBT *Drosophila* embryos, in the absence of external damage, and assess the consequences of this activity during development from larvae to adult stage, we employed a previously generated *dpolη^Exc2.15^*null mutant to obtain dPolη maternally-depleted progeny (*40*). Eggs laid by females flies homozygous for the *dPolη^Exc2.15^*mutation, are maternally-deprived of dPolη because in the pre-MBT embryos transcription is naturally shut off and is only re-established after MBT by transcription of the paternal gene provided by the sperm of wild-type flies (Figure 3A). By consequence, phenotypes observed in this experimental setting can be attributed to dPolη absence during the pre-blastoderm cleavage stages. dPolη maternally-depleted flies were obtained by crossing either *dPolη^Exc2.15^/ dPolη^Exc2.15^* (homozygous ^−/−^), or *dPolη^Exc2.15^*/+ (heterozygous ^+/−^, as a control) females with wild-type male flies (Figure 4A). Development was monitored from early embryos to adult stage and compared to that of both wild-type and maternally-provided flies (obtained by crossing wild-type males with *dPolη^Exc2.15/+^* flies as explained above). Pre-blastoderm *dPolη* maternally-depleted embryos showed altered chromatin features, suggesting defects in chromosome segregation (Figure 4B). Consistent with these observations, and in contrast with previously published data (*40*)*, dPolη* maternally-deprived embryos exhibited higher mortality and reduced hatching rate compared to maternally-provided flies embryos whose hatching rate was similar to that of the wild-type (Figure 4C). We observed a very similar phenotype in compound mutant flies, bearing two different *dPolη* alleles (*dPolη^Exc2.15^* and *dPolη^12^*), confirming that the observed reduced hatching rate is specifically due to dPolη absence. Altogether, these results suggest that there is no dPolη dosage effect. These phenotypes are entirely consistent with a similar phenotype very recently reported in TLS-deficient *C. elegans pcna^K164R^* mutant embryos (*42*). No significant difference in the survival rate was detected from larvae to adult stage in *dPolη*^EXC215^ crosses (Figure 4D). In conclusion, these results indicate that dPolη absence before MBT affects embryo’s survival from embryo to larva stage.

**Figure 4.**
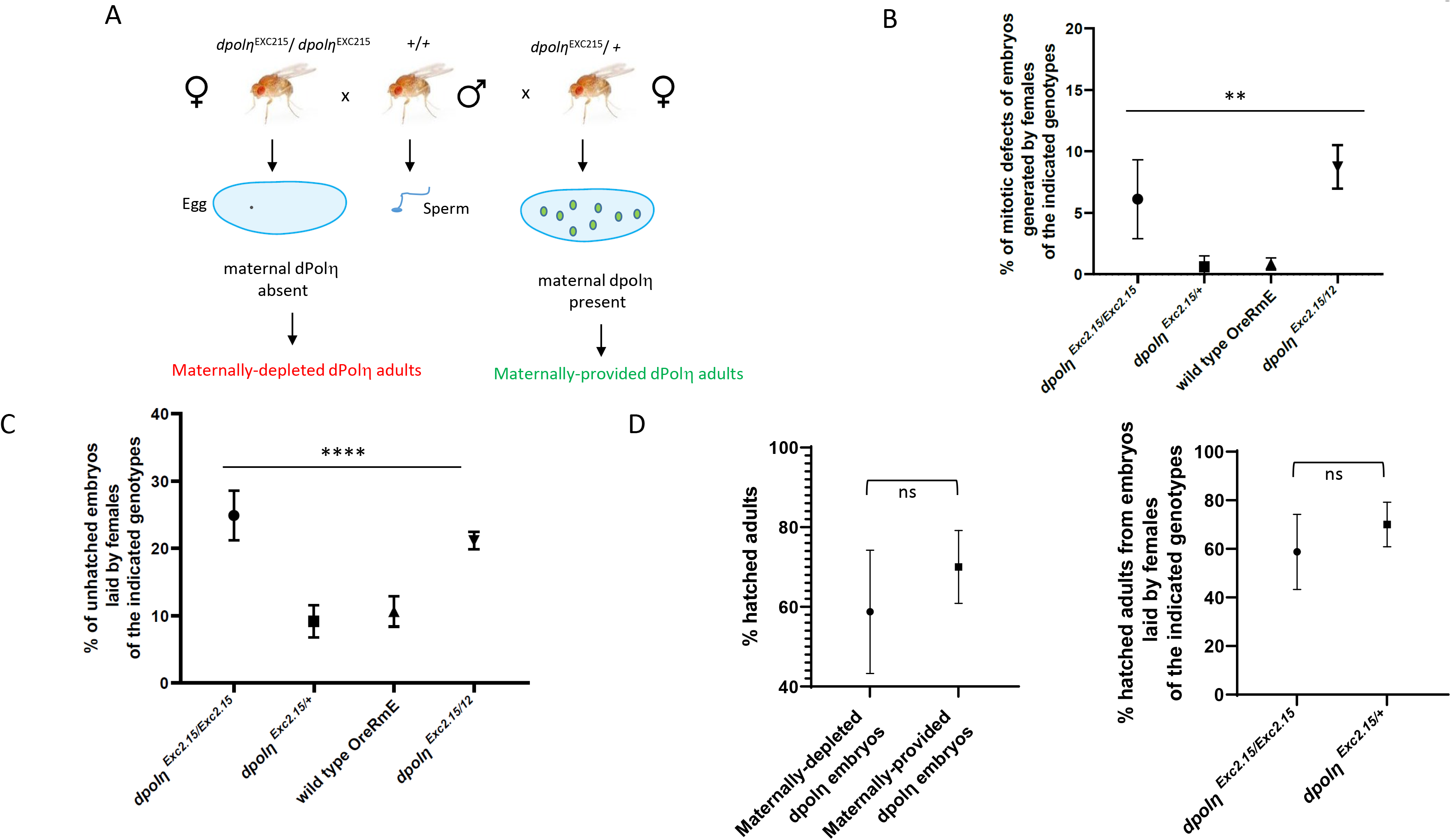
Absence of maternal dPol decreases embryo’s hatching rate. (A) Drawing of the experimental plan employed to evaluate the importance of dPolη during *Drosophila* early embryogenesis. In this experimental set up, dPolη is provided only after MBT upon zygotic gene activation. (B)Quantification of embryos displaying mitotic defects (n=2). Data are presented as means ± SD. Means were compared using analysis of one-way ANOVA. (C) Statistical analysis of embryo-to-larva hatching rate of *Drosophila* embryos laid by mothers bearing the indicated *dPolη* genotypes. Embryos were collected over 12 hours and incubated at 25 °C for 2 days before calculating hatching rate. Means and standard deviation of unhatched embryos are expressed as error bars. Means were compared using analysis of one-way ANOVA followed by unpaired Student’s t test for the first panel (from left to right) and Fisher’s exact test for the second panel (n=3). (D) Statistical analysis of larva-to-adult transition hatching rate in the absence (maternally depleted) or presence (maternally provided) of maternal dPolη (left panel), and comparison between two different *dpol* mutants (right panel). Data are presented as means ± SD. Means were compared with unpaired Student’s t test (n=3).

### Genome-wide analysis of *dPolη* maternally-depleted flies

To determine dPolη contribution to mutagenesis during the *Drosophila* cleavage stages, we analyzed SNVs, short insertions and deletions (Indels) by Whole Genome Sequencing (WGS) in the genome of single flies obtained by crossing females either *dPolη^Exc2.15^*/ *dPolη^Exc2.15^* (homozygous ^−/−^), or *dPolη^Exc2.15^*/+ (heterozygous ^+/−^), with the same wild-type Oregon-R male fly homogenized through serial individual backcrosses for nine generations, so to obtain genotypically identical progeny (Supplementary Figure S4B). Due to the error-prone nature of TLS Polη, especially on undamaged templates, it is expected that flies obtained from maternally-depleted *dpolη* embryos must display less mutations than either flies that contain maternally-deposited dPolη or wild-type flies. However, following alignment against the reference genome (dm6) and general genome analysis, we observed that the global number of SNVs and Indels was not significantly reduced (Figure 5A-B), consistent with previous observations in *C. elegans* (*43*). This result can be interpreted as functional redundancy amongst Y-family TLS pols (*9*, *44*). Notwithstanding, analysis of SNVs distribution on specific chromosomes revealed a significant SNV depletion on chromosome 3 of maternally-depleted flies compared to maternally provided flies (Supplementary Figure S4A-B and Figure 5C), in particular within a cluster of Responder (Rsp) satellite DNA repeat within the pericentromeric heterochromatin of chromosome 3L (Figure 5D). The SNVs difference in this region accounts for 100-fold decrease in the mutagenesis rate in the maternally-depleted flies compared to the maternally-provided flies, consistent with an error-prone activity of dPolη. A difference was also observed in the pericentromeric region of the right part of the same chromosome (3R) including a shift far from the centromere in the maternally-depleted flies (Figure 5D), while no gross variations were observed on other chromosomes, except some significant variations on chromosome 4, X and Y (Supplementary Figure S4D-F). Analysis of the mutation spectrum on chromosomes 3R and 3L (Figure 5C) revealed a predominant reduction of C>T, T>C transitions and T>A transversions. This is in line with the observation that unlike yeast and humans, dPolη misincorporates G opposite T template leading to T>C transitions (*45*).

**Figure 5.**
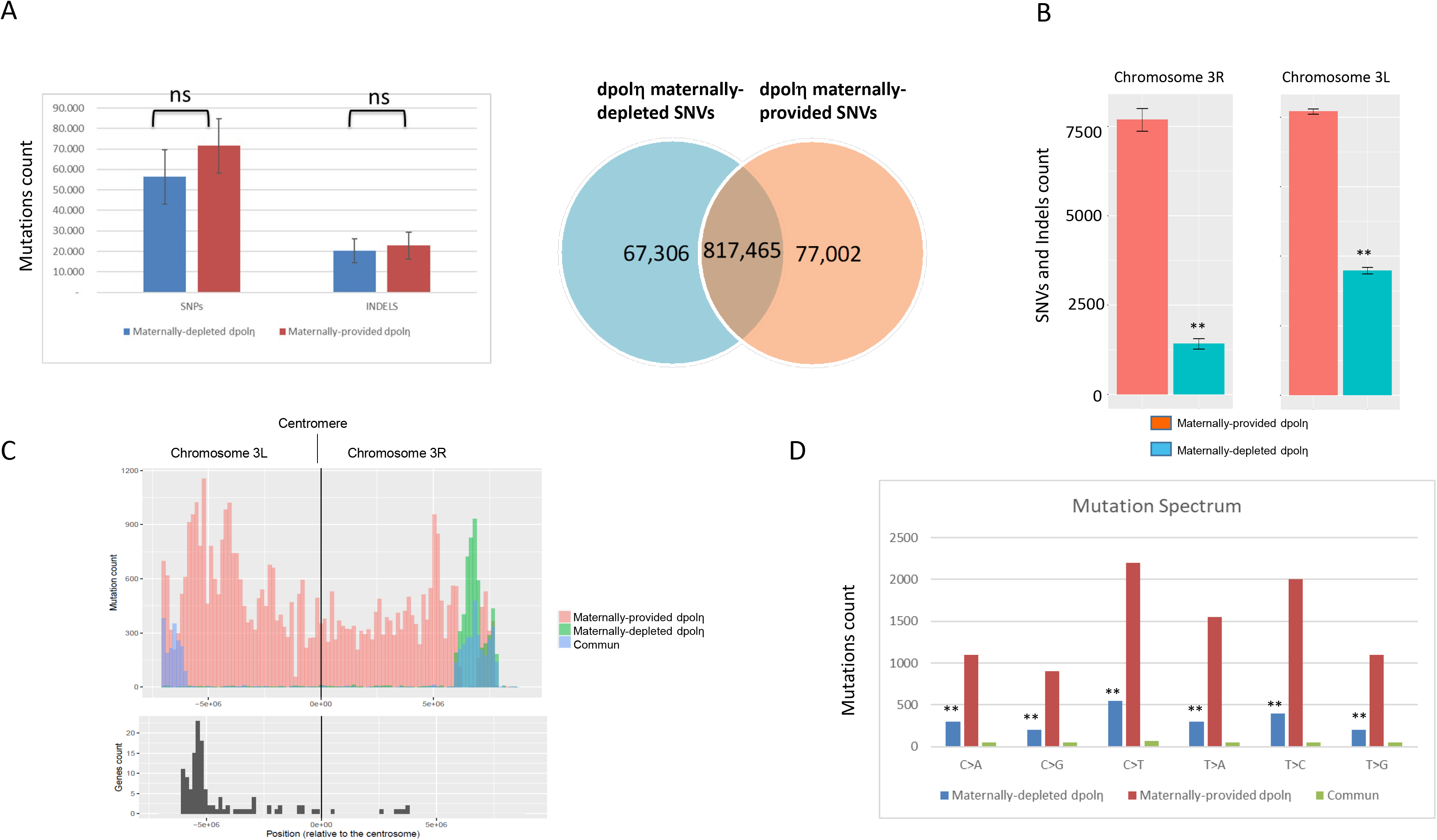
Chromosome-specific variation in SNVs in absence of maternal dPolη. (A, left panel) SNVs and Indels abundace in WGS sequenced adult flies either maternally depleted or maternally provided of dPolη. Right panel, Diagram displaying how many counted SNVs on the indicated fly genomes are mutual or distinct. The graph represents the average of two sequenced flies per condition. Data are presented as means ± SD. No statistical significant difference was detected for either SNVs or Indels (unpaired Student’s t test). (B) Chromosome 3L (left panel) and 3R (right panel) SNVs rate of maternally-provided (red bars) or maternally-depleted (green bars) *dpol* males adult flies. SNVs means of two independent WGS analyses are expressed as counts in the columns. Data are presented as means ± SD. Means were compared using unpaired Student’s t test (n=2). (C) SNVs graphic representation (mutation count) in the pericentromeric region of either the left portion (3L) or right portion (3R) of chromosome 3. SNVs found only in either maternally-depleted or maternally-provided dPolη progeny are shown respectively in orange or green, while common SNVs are illustrated in blue. Lower panel, graphic representation of genes density (genes count) along the indicated chromosome regions (n=2). (D) Mutation spectrum on chromosome 3 where x-axis indicates types of nucleotide variants and y-axis the quantity of counted SNVs. Unique polymorphism in either maternally-depleted or partially maternally-provided dPolη progeny are separately represented in blue or in red, while mutual variants are displayed in green. Means were compared using unpaired Student’s t test (n=2).

Because approximately two third of the genes located on 3L pericentromeric heterochromatin are required for developmental viability and/or adult fertility (*46*) we evaluated the predicted effects of mutations in either maternally-depleted or maternally-provided adults by attributing variant effect predictor (VEP) score to each variation. VEP score determines the effect of variants (SNVs, insertions, deletions, CNVs or structural variants) on genes, transcripts, and protein sequence, as well as regulatory regions. In both maternally-depleted and maternally-provided flies genomes, the majority of variants present a modifier score more enriched in the maternally-depleted mutant on the pericentromeric region of chromosome 3L and 3R (Supplementary Figure S5A). Most variants in this category affect intron splicing or noncoding regions (intergenic variants, Supplementary Figure S5A-B). However, once the VEP score modifier removed, the most recurrent SNVs presented a moderate score or a low score, (Figure 5C-D). Scoring the consequences of these variants shows that they lead to missense mutations in coding genes in the maternally-deprived mutant. This category of variants changes the genetic code, which may potentially alter the function of a protein.

Taken altogether, these data show that *dPolη* maternally-depleted adults are characterized by decreased mutations on specific chromosomes regions that may depend upon dPolη for efficient replication during the very fast cleavage stages and containing genes important from embryos viability (*46*).

## Discussion

### High mutagenesis in the very early *Xenopus* embryo

In this work, we have provided evidence for the occurrence of a surprisingly high mutation rate in the very early, pre-MBT *Xenopus* embryo. Mutation rate was estimated to be in the range of 10^−3^ and corresponds to 0.8 mutations per cell cycle, a value very close to that observed in the human germline (0.4-1.2) (*47*) but slightly lower than that estimated for pre-implantation human embryos (2.8) (*48*). This mutation rate, which is within the range observed for Y-family TLS Pols, was greatly reduced upon expression of either the TLS-deficient Rad18^C28F^ or the Rad18^C28FC207F^ mutant. The corresponding mutation spectrum is also consistent with the mutagenesis spectrum of TLS Pols on undamaged DNA templates. Collectively, these findings indicate that in pre-MBT *Xenopus* embryos TLS strongly contributes to the observed mutagenesis.

The residual mutations observed in the Rad18^C28F^ mutant includes C>T transitions and C>A transversions. These mutations, that were recently reported to be also predominant in early human embryos (*48*), can be a consequence of either ribonucleotides incorporation or generated as a result of cytosine deamination into uracil, which is then turned into a thymine upon replication. The concentration of ribonucleotides exceeds of about 1000-fold that of deoxyribonucleotides, and we have observed a high level of ribonucleotides incorporation during DNA synthesis in *Xenopus* egg extracts that depends upon the DNA-to-cytoplasmic ratio (Supplementary Figure S2E). This may depend upon TLS activity, in line with evidence demonstrating ribonucleotides incorporation *in vitro* by human Polη (*49*, *50*). Because ribonucleotides slow down DNA replication, constitutive TLS activation facilitates their bypass, a strategy that the *Xenopus* embryos may have evolved to cope with a highly contracted cell cycle. Notwithstanding, it cannot be excluded that these mutations might be also a consequence of unproofed errors of replicative DNA Pols.

Unexpectedly, we have also observed large deletions in the *lacZ* gene recovered from pre-MBT embryos. These rearrangements are unlikely to be an artifact of the plasmid assay, since they were not detected neither in plasmids isolated from embryos co-injected with Rad18^WT^, nor from UV-irradiated embryos. In this respect, genomic deletions have been observed in the *S* subgenome of adult *Xenopus laevis* (*51*) as well as in *Drosophila melanogaster* (*52*), suggesting that such genomic rearrangements might be generated naturally during evolution in these organisms. Although Y-family TLS Pols can generate deletions, their extent is rather small (1-3 bp), implying other mechanisms such as replication fork instability which is a common feature of DNA damage checkpoints inefficiency (*5*, *53*, *54*). Replication fork collapse can also happen when repriming and template switch are inefficient, however we have found no evidence for repriming nor for template switching in the context of *Xenopus* early embryogenesis (*10*) (and our unpublished observations). Replication fork collapse can be a consequence of suboptimal TLS activity due to limiting Rad18 levels (*10*). Consistent with this interpretation, *C. elegans* strains with reduced TLS function accumulate spontaneous genomic deletions as a result of double strand breaks at forks arrested by endogenous DNA lesions (*43*). Notwithstanding, it cannot be excluded that these rearrangements are the result of rare intermediates, Polη-dependent, that turned into deletion upon transformation into *E.coli*. Rad18^WT^ overexpression would reduce fork stalling by boosting both TLS and HDR, suppress NHEJ toxic effect at collapsed replication forks and therefore reduce deletions. This scenario is in line with evidence showing that Rad18 has a negative effect on NHEJ (*55*) and that NHEJ is predominant over HDR in the early *Xenopus* embryo (*56*). Externally applied DNA damage may stimulate repriming and/or template switch by so far unclear mechanisms, thus facilitating replication fork restart, and suppressing replication fork collapse.

### Functional conservation of constitutive TLS in the early embryogenesis of fast cleaving embryos

Similar to what observed in *Xenopus* (*10*), we have provided evidence for both developmental regulation of PCNA^mUb^ in *Drosophila* early embryogenesis, and TLS activity, suggesting that this process is also conserved in invertebrates. Because a *Drosophila* Rad18 ortholog in *Drosophila* could not be identified, it has not been possible to analyze mutagenesis in a complete Y-family TLS-free context.

Detailed genome-wide analysis of SNVs in maternally-depleted *dPolη* adults revealed a strong SNVs reduction in the pericentromeric region of chromosome 3, as well as SNVs depletion on the Y chromosome and the pericentromeric region of chromosome X. It is currently unclear why dPolη maternal depletion mainly affects SNVs abundance on chromosome 3. This is the largest *Drosophila* chromosome which includes the largest cluster of 120 bp *Responder* DNA repeats of α-satellite DNA within the pericentromeric heterochromatin (*57*). Such DNA sequences form secondary structures that constitute a challenge for a canonical replication fork, and Polη, and not Polι, has been previously shown to be important to replicate unusual DNA sequences in somatic cells (*58*, *59*). Because *Drosophila* lacks Polκ, Polη may be essential to assist the replisome in the replication of heterochromatin, in particular on chromosome 3 that contains the largest block of *Responder* DNA repeats. This interpretation is consistent with SNVs depletion observed on chromosome Y which is highly heterochromatic and with the mutation spectrum that corresponds to the reported incorporation errors of Polη on undamaged templates (*33*, *35*, *45*). Either replicative polymerases, or dRev1 may compensate for dPolη absence, although less efficiently. Due to inefficiency of the replication checkpoint in the *Drosophila* pre-blastoderm syncytium, embryos may accumulate chromosome abnormalities and undergo apoptosis at MBT (*5*), thus explaining the reduced hatching rate and chromosome abnormalities observed in maternally-depleted *dpol η* embryos. A caveat of this interpretation is that mapping to highly repetitive genomic regions is not very accurate. However, we do not have any indication that this contributed to the difference in SNVs identified on 3L and 3R chromosome arms between the maternally-depleted and maternally-provided dPol η flies.

The 3L chromosome region also contain a set of genes involved in development and viability. A great majority of SNVs in this region are predicted to generate mutations with low or moderate impact on genes functions. Hence, it cannot excluded that the phenotype observed in the *dpolη* homozygous flies may also be a consequence of mutations in essential genes.

### Consequences of a high mutagenic rate in early embryogenesis: good or bad?

The occurrence of a high mutation rate in early developing embryos of fast cleaving organisms is rather surprising but somehow not completely unexpected since these embryos are characterized by a highly contracted cell cycle that does not leave enough time to allow quality control (*1*). In this situation, the toll to pay is an increased risk of mutagenesis and genomic instability, as we have reported in this work. Several reports have highlighted the occurrence of genomic instability and mutations in early embryos (*1*) for review), which is apparently compatible with normal development (*63*). These observations suggest that active protection mechanisms must be operating to reduce the mutation load. Consistent with this possibility, mutagenesis dropped to background levels when *Xenopus* embryos were injected with a high dose of DNA, which mimics a pre-MBT stage (*6*). At MBT, cell cycle extension, activation of the DNA damage checkpoint and apoptosis would ensure repair of errors introduced during the cleavage stages, thereby limiting the propagation of cells having gross chromosomal alterations, and explaining both the low level of developmental defects and embryonic mortality (*5*, *60*–*62*). In this work, we have unveiled that in *Xenopus*, Rad18 has a protective function through an error-free HDR activity that reduces its TLS mutagenic activity. In addition, we have shown that the MMR pathway also contributes to reduce mutagenesis in the pre-MBT embryo. However, we have shown that in *Drosophila*, mutations generated in the pre-blastoderm embryo are inherited in the adult, suggesting that protection mechanisms against genomic instability are not very stringent.

Introduction of random mutations, generated by so far unclear mechanisms, has been also recently observed in human embryos (*48*). In addition, pre-implantation human embryos display genomic instability characterized by gross chromosomal rearrangements that can lead to cleavage arrest at the 2-4 cell stage (*63*). Introduction of random mutations may constitute an unexpected and novel source of genetic variation contributing to genome evolution that may be advantageous for the adaptation of the species, but at the same time might be dangerous for life. For example, an overall high mutation rate may be important for pseudogenization, a process that silences the expression of pseudogenes (*64*) and be also important to adaptation to a new environment. A recent study identified several genes located on the *Drosophila* 3L chromosome involved in adaptation (*65*). We have observed that Polη mutagenic activity may be important to maintain the stability of centromeric DNA sequences in the *Drosophila* pre-blastoderm embryo, thus being good for life. Genome-wide association studies have implicated hundreds of thousands of single-nucleotide polymorphisms (SNPs) in human diseases and traits (*66*). In the future, it will be important to explore which is the level of DNA damage inherited in the post-MBT embryo, and its contribution to the polymorphisms that characterize each individual.

## Acknowledgments

We wish to thank Marcel Méchali for technical advises, J-S Hoffmann and SE Kearsey for critical reading of the manuscript, B Rondinelli, J Sale, A-M Martinez, A Goriely for useful discussions and H-M Bourbon for help with sequence alignments. This project was supported by ANR (ANR-12-BSV2-0022) and MSD Avenir grants to D.M. E.L.F. was supported by a 3-year PhD fellowship from the “Ligue contre le Cancer” and 1-year fellowship from “Fondation ARC contre le cancer”.

## Competing Interests

The authors declare that they have no competing interests

## Author contributions

Conceptualization, D.M., E.L.F.; Methodology, D.M., E.L.F., I.B.; Investigation, E.L.F., I.B.; Formal Analysis, C.S., S.Z., C.M.D., C.L.; Writing, E.L.F., D.M.; Funding Acquisition, D.M.; Supervision, D.M., I.B.; Visualization, D.M. and E.L.F.; Resources, D.M. and I.B.

## Data repository

All data needed to evaluate the conclusions in the paper are present in the paper and/or the Supplementary Materials. WGS data have been submitted to Gene Expression Omnibus (GEO, https://www.ncbi.nlm.nih.gov/geo/; ID GSE161335).

---

**Figure S1.**
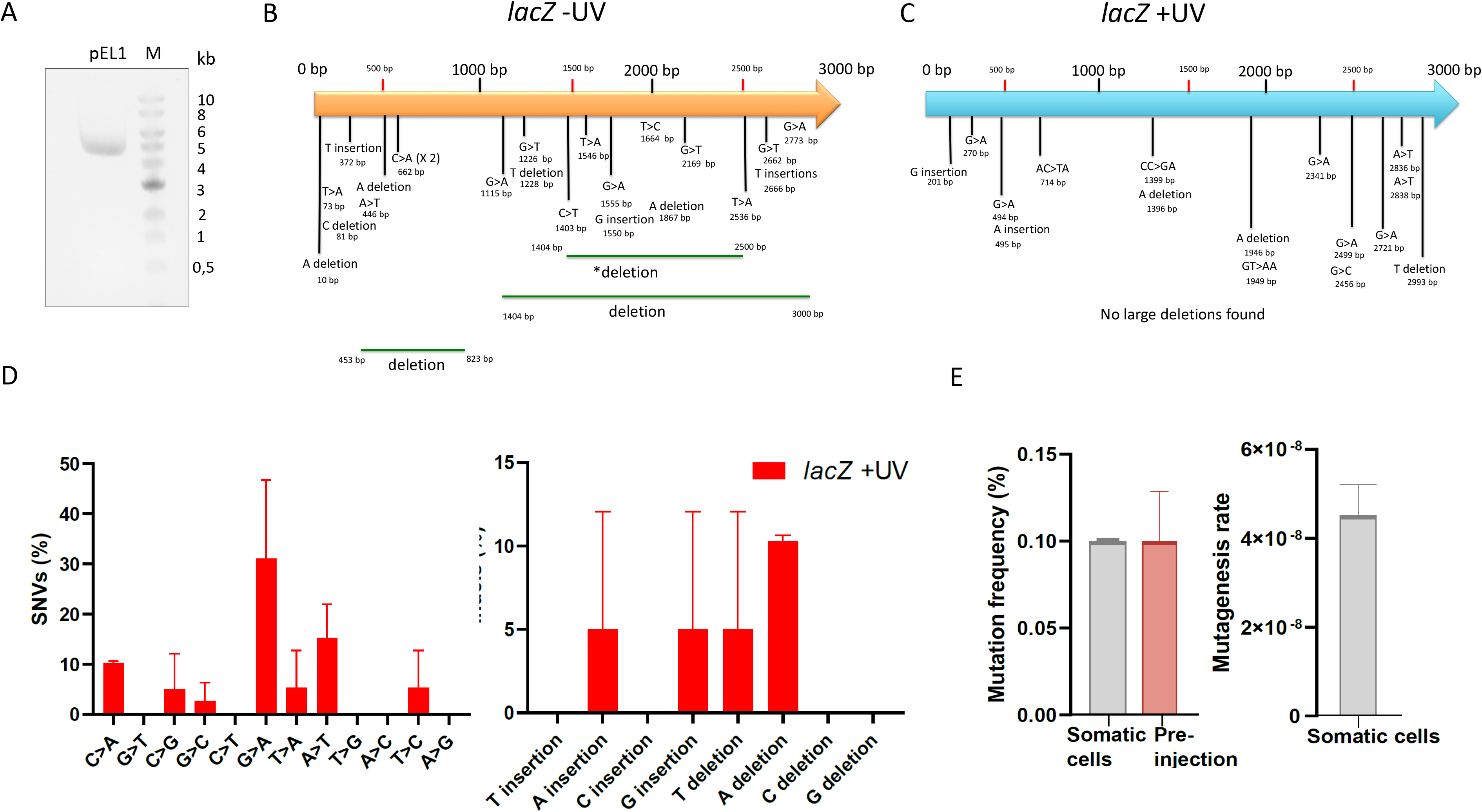
Mutation maps and SNVs in early *Xenopus* embryos. (A) Agarose gel electrophoresis of *lacZ-*containing plasmid (pEL1). M indicates DNA ladder molecular weight marker. (B-C) Mutations map (SNVs, insertions, deletions) on the *lacZ* gene recovered from pre-MBT embryos irradiated (+UV) or not (*lacZ*) with UV-C (n=3). (D) Mutation spectra of UV-irradiated *lacZ* gene recovered from *Xenopus* pre-MBT embryos after Sanger sequencing. Data are presented as means ± SD (n=2). (E) Mutation rate (left panel) and frequency (right panel) of cultured somatic A6 *Xenopus* cells. Mutagenesis rate is expressed as mutations per base pair/locus per generation (see Materials and Methods), normalized to the pre-injection background values. Data are presented as means ± SD (n=2).

**Figure S2.**
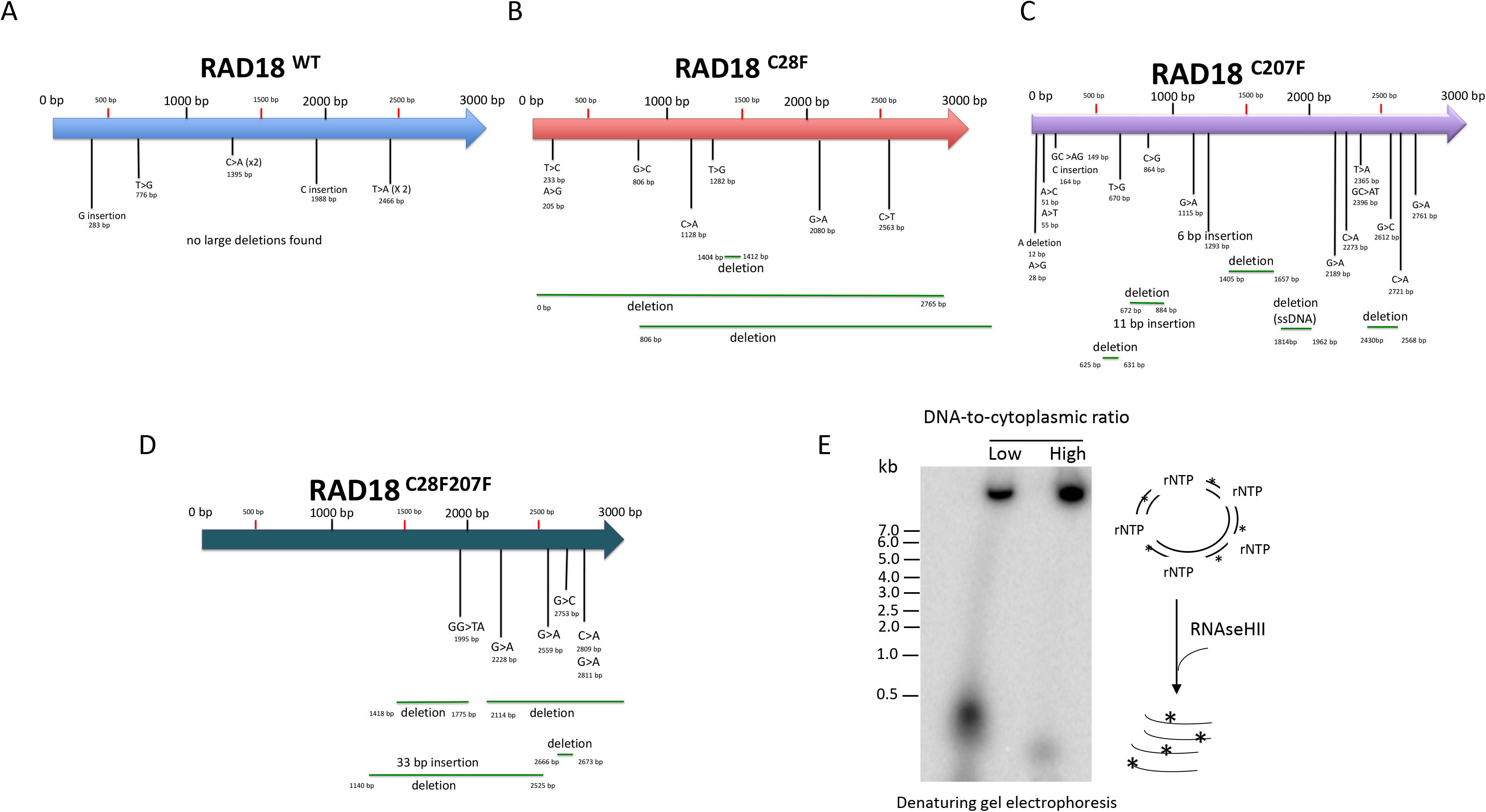
Mutations map of *lac* Z co-injected with RAD18 mRNAs. (A-D) Mutations map of the *lacZ* gene recovered from pre-MBT embryos co-injected with the indicated Rad18 forms (n=3). (E) High level of ribonucleotides incorporation during DNA synthesis at low DNA-to-cytoplasmic ratio. Autoradiograph of radioactive-labeled M13 ssDNA replicated in egg extracts at different doses: low (0,66 ng/μL^−1^) for pre-MBT-like or high DNA-to-cytoplasmic ratio (6,6 ng/ μL^−1^) for post-MBT-like dose, treated with NaOH and separated by denaturing urea-gel electrophoresis. Products were compared with DNA ladder of know size (n=2).

**Figure S3.**
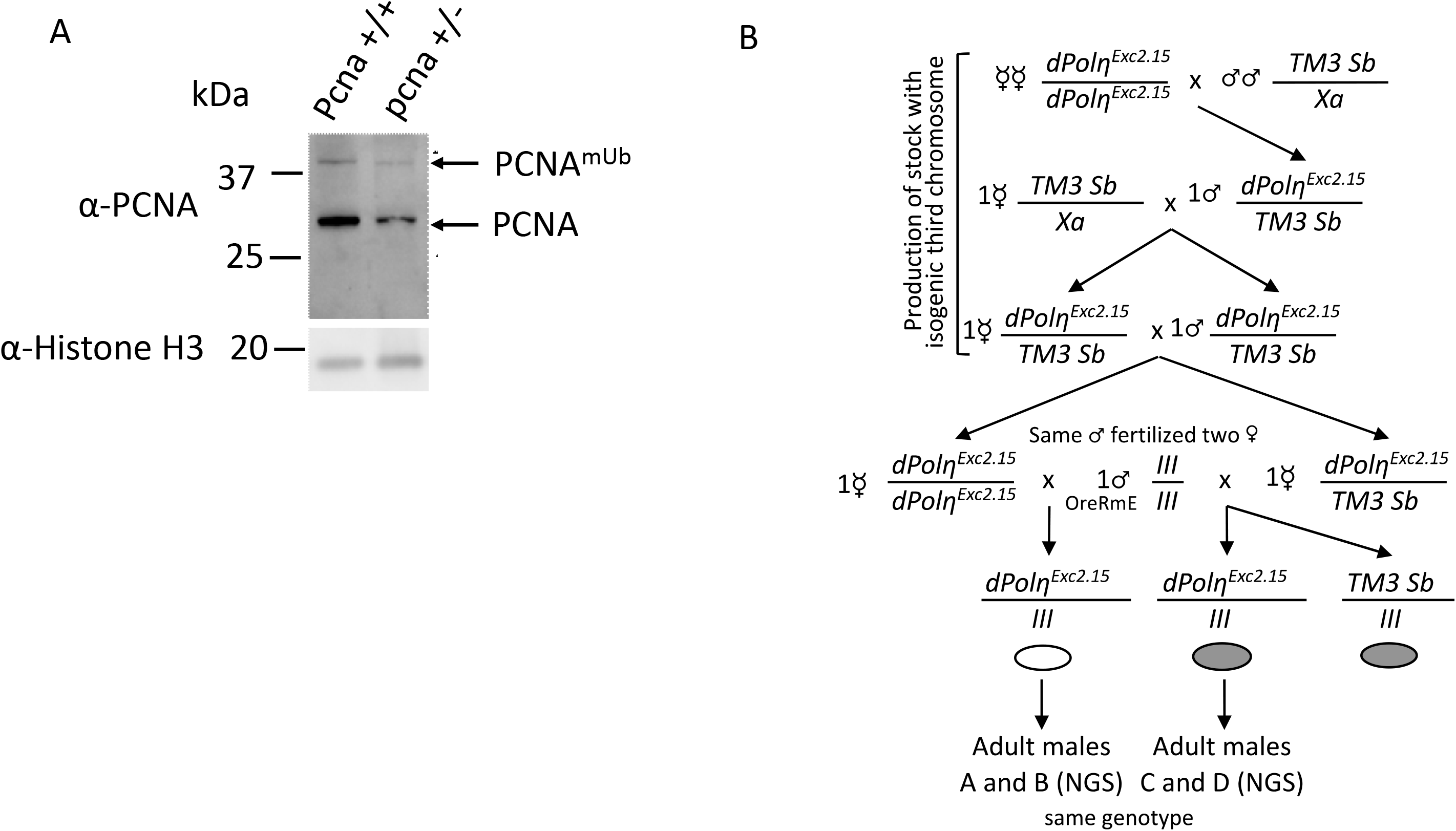
PCNA detection in *Drosophila* and genetic crosses used to generate maternally-depleted *dpolη* flies. (A) Western blot of total protein extracts obtained from flies of the indicated genotype with the PC10 antibody (n=2). (B) Genetic crosses set up to generate either maternally-depleted or maternally-provided dPolη adult flies. The first two crosses generated a balanced stock over TM3 marked with Sb with a isogenic 3rd chromosome carrying the *dPolη^Exc2.15^* mutation. From this stock, one homozygous (left) and one hetreozygous (right) virgin females were collected and mated with the same wild type male, to produce embryos which were respectively depleted or not of maternal deposited dPolη. Then, genotypically identical adult males, devoid of the Sb marker, and containing the isogenic 3rd chromosome carrying the *dPolη^Exc2.15^* mutation, were collected for WGS processing and analysis.

**Figure S4.**
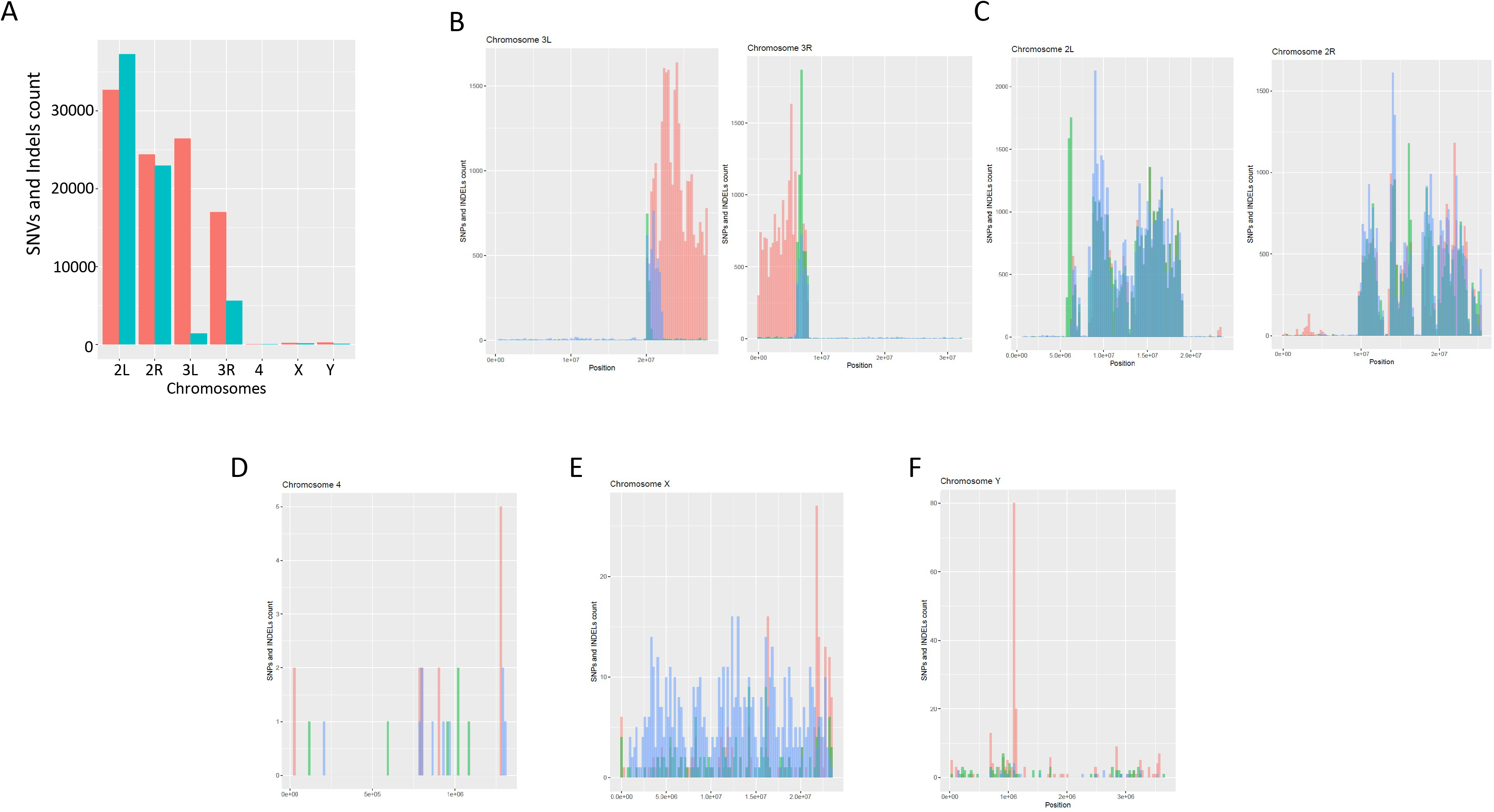
Distribution of mutations in the absence of maternal dpol across chromosomes. (A) SNVs and Indels count on the indicated *Drosophila melanogaster* chromosomes. (B-F) Mean of SNVs and Indels distribution on the indicated *Drosophila* chromosomes (n=2). The x-axis describes the position of counted variants on the chromosome and y-axis displays how many variants were counted on that position. Means were compared with an unpaired-two tailed Student’s t test.

**Figure S5.**
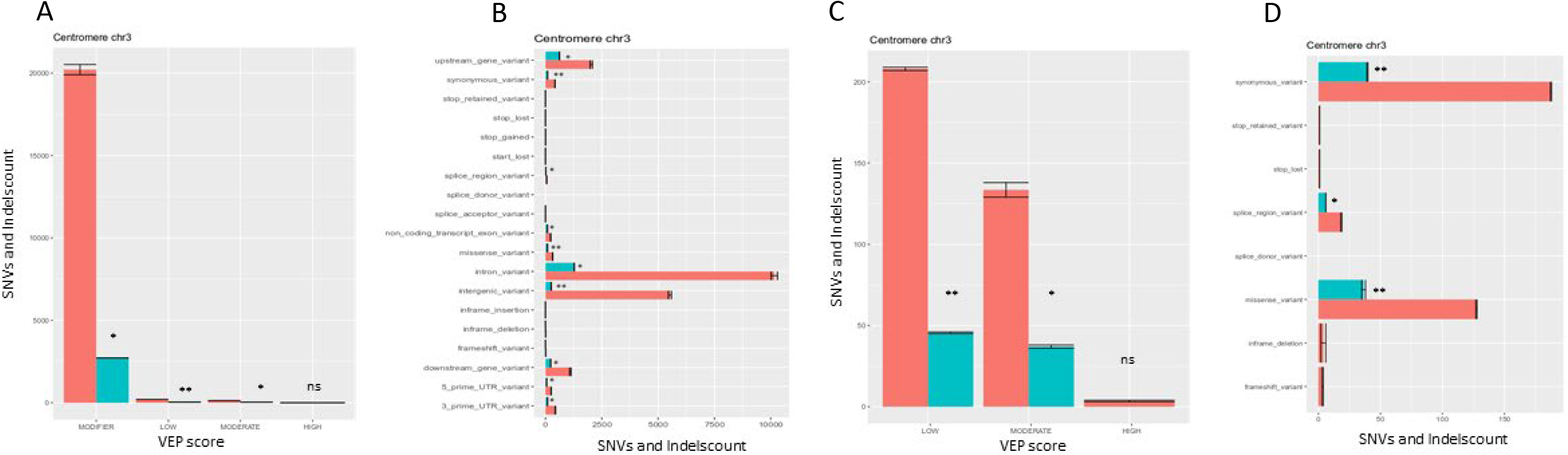
Impact of dPolη maternal depletion on gene function. (A) Diagram classifying the impact of mutations found on chromosome 3L (left panel) and chromosome 3R (right panel). Mutations are classified according to their effect by attribution of a VEP score. These panels describe how many SNVs (y-axis) have high, low, moderate or modifier effect (x-axis) in each sequenced fly. Most mutations generated in maternally-provided dPolη flies do not particularly affect genes, transcripts, protein sequence, or regulatory regions (n=2). (B) In this plot, types of variants with modifier VEP score are listed on the y-axis and the counts for the type of polymorphisms are shown on the x-axis. Most of mutations are slightly altering introns or intergenic regions (n=2). (C) In this panel, modifier VEP scores have been discarded to focus on the consequences of other scored variants. Most polymorphisms have a moderate effect on altered gene (n=2). (D) Evaluation of mutation impact upon discarding variants with modifier score. In this graph, types of variants with other scored are illustrated on the y-axis and the counts for each type of polymorphisms are shown on the x-axis. Most of SNVs give raise to missense variants (n=2). Data are presented as means ± SD. Means were compared with an unpaired Student’s t test.

## Notes

https://www.ncbi.nlm.nih.gov/geo/

### Competing Interest Statement

The authors have declared no competing interest.

